# An extension of the *studyforrest* dataset for vision research

**DOI:** 10.1101/046573

**Authors:** Ayan Sengupta, Falko R. Kaule, J. Swaroop Guntupalli, Michael B. Hoffmann, Christian Häusler, Jörg Stadler, Michael Hanke

## Abstract

The *studyforrest* (http://studyforrest.org) dataset is likely the largest neuroimaging dataset on natural language and story processing publicly available today. In this article, along with a companion publication, we present an update of this dataset that extends its scope to vision and multi-sensory research. 15 participants of the original cohort volunteered for a series of additional studies: a clinical examination of visual function, a standard retinotopic mapping procedure, and a localization of higher visual areas — such as the fusiform face area. The combination of this update, the previous data releases for the dataset, and the companion publication, which includes neuroimaging and eye tracking data from natural stimulation with a motion picture, form an extremely versatile and comprehensive resource for brain imaging research — with almost six hours of functional neuroimaging data across five different stimulation paradigms for each participant. Furthermore, we describe employed paradigms and present results that document the quality of the data for the purpose of characterising major properties of participants’ visual processing stream.

## Background & Summary

The *studyforrest* dataset^1^, with its combination of functional magnetic resonance imaging (fMRI) data from prolonged natural auditory stimulation and a diverse set of structural brain scans, represents a versatile resource for brain imaging research with a focus on information processing under real-life like conditions. The dataset has, so far, been used to study the role of the insula in dynamic emotional experiences^2^, modeling of shared blood oxygenation level dependent (BOLD) response patterns across brains^3^, and to decode input audio power-spectrum profiles from fMRI^4^. The dataset has subsequently been extended twice, first with additional fMRI data from stimulation with music from various genres^5^ and secondly with a description of the movie stimulus structure with respect to portrayed emotions^6^. However, despite providing three hours of functional imaging data per participant, experimental paradigms exclusively involved auditory stimulation, thereby representing a substantial limitation regarding the aim to aid the study of real-life cognition — which normally involves multi-sensory input. With this further extension of the dataset presented here and in a companion publication^7^, we are now substantially expanding the scope of research topics that can be addressed with this resource into the domain of vision and multi-sensory research.

This extension is twofold. While the companion publication^7^ describes an audio-visual movie dataset with simultaneously acquired fMRI, cardiac/respiratory traces, and eye gaze trajectories, the present article focuses on data records and exams related to a basic characterization of the functional architecture of the visual processing stream of all participants — namely retinotopic organization and the localization of particular higher-level visual areas. The intended purpose of these data is to perform brain area segmentation or annotation using common paradigms and procedures in order to to study the functional properties of areas derived from these standard definitions in situations of real-life like complexity. Moreover, knowledge about the specific spatial organization of visual areas in individual brains aids studies of the functional coupling between areas, and it also facilitates the formulation and evaluation of network models of visual information processing in the context of the *studyforrest* dataset.

The contributions of this study comprise three components: 1) results of a clinical eye examination for subjective measurements of visual function for all participants to document potential impairments of the visual system that may impact brain function, even beyond the particular properties relevant to the employed experimental paradigms; 2) raw data data for a standard retinotopic mapping paradigm and a six-category block-design localizer paradigm for higher visual areas, such as the fusiform face area (FFA)^8^, the parahippocampal place area (PPA)^9^, the occipital face area^10^, the extrastriate body area (EBA)^11^, and the lateral occipital complex (LOC)^12^; and 3) validation analyses providing volumetric angle maps for retinotopy data and ROI masks for visual areas. While the first two components are factual, the third component is based on a largely arbitrary selection of analysis tools and procedures. No claim is made that the chosen methods are superior to any alternative, but the results are shared to document the plausibility of the results and to facilitate follow-up studies that do not require any particular method to analyze and interpret these data.

## Methods

### Participants

Fifteen right-handed participants (mean age 29.4 years, range 21-39, 6 females) volunteered for this study. All of them had participated in both previous studies of the *studyforrest* project^1,5^. The native language of all participants was German. The integrity of their visual function was assessed at the Visual Processing Laboratory, Ophthalmic Department, Otto-von-Guericke University, Magdeburg, Germany as specified below. Participants were fully instructed about the purpose of the study and received monetary compensation. They signed an informed consent for public sharing of all obtained data in anonymized form. This study was approved by the Ethics Committee of the Otto-von-Guericke University (approval reference 37/13).

### Subjective measurements of visual function

To test whether the study participants had normal visual function and to detect critical reductions of visual function, two important measures were determined: (1) visual acuity to identify dysfunction of high resolution vision and (2) visual field sensitivity to localize visual field defects. For each participant, these measurements were performed for each eye separately — if necessary with refractive correction. (1) Normal decimal visual acuity (>=1.0) was obtained for each eye of each participant. (2) Visual field sensitivities were determined with static threshold perimetry (standard static white-on-white perimetry, program: dG2, dynamic strategy; OCTOPUS Perimeter 101, Haag-Streit, Koeniz, Switzerland) at 59 visual field locations in the central visual field (30° radius) *i*.*e*., covering the part of the visual field that was stimulated during the MRI scans. In all, except for two participants, visual field sensitivities were normal for each eye (MD (mean defect) dB<2.0 & >-2.0; LV (loss variance) dB^2^ < 6) — indicating the absence of visual field defects. Visual field sensitivities for participant 2 (right eye) and participant 4 (both eyes) were slightly lower than normal but not indicative of a distinct visual field defect.

### Functional MRI acquisition setup

For all of the fMRI acquisitions described in the paper, the following parameters were used: T2*-weighted echo-planar images (gradient-echo, 2 s repetition time (TR), 30 ms echo time, 90 ° flip angle, 1943 Hz/px bandwidth, parallel acquisition with sensitivity encoding (SENSE) reduction factor 2) were acquired during stimulation using a whole-body 3Tesla Philips Achieva dStream MRI scanner equipped with a 32 channel head coil. 35 axial slices (thickness 3.0 mm) with 80x80 voxels (3.0x3.0 mm) of in-plane resolution, 240 mm field-of-view (FoV), anterior-to-posterior phase encoding direction) with a 10% inter-slice gap were recorded in ascending order — practically covering the whole brain. Philips’ “SmartExam” was used to automatically position slices in AC-PC orientation such that the topmost slice was located at the superior edge of the brain. This automatic slice positioning procedure was identical to the one used for scans reported in the companion article^13^ and yielded a congruent geometry across all paradigms.

### Physiological recordings

Pulse oximetry and recording of the respiratory trace were performed simultaneously with all fMRI data acquisitions using the built-in equipment of the MR scanner. Although the measurement setup yielded time series with an apparent sampling rate of 500 Hz, the effective sampling rate was limited to 100 Hz.

### Stimulation setup

Visual stimuli were presented on a rear-projection screen inside the bore of the magnet using an LCD projector (JVC DLA RS66E, JVC Ltd., light transmission reduced to 13.7% with a gray filter) connected to the stimulus computer via a DVI extender system (Gefen EXT-DVI-142DLN with EXT-DVI-FM1000). The screen dimensions were 26.5cmx21.2cm at a resolution of 1280 x 1024 px with a 60 Hz video refresh rate. The binocular stimulation were presented to the participants through a front-reflective mirror mounted on top of the head coil at a viewing distance of 63 cm. Stimulation was implemented with PsychoPy v1.79 (with an early version of the `MovieStim2` component later to be publicly released with PsychoPy v1.81)^14^ on the (Neuro)Debian operating system^15^. Participant responses were collected by a two-button keypad and was also logged on the stimulus computer.

### Retinotopic Mapping

#### Stimulus

Similar to previous studies^16,17^, traveling wave stimuli were designed to encode visual field representations in the brain using temporal activation patterns^18^. Expanding/contracting rings and clockwise/counter-clockwise wedges (see Figure 3A) consisting of flickering radial checkerboards (flickering frequency of 5 Hz) were displayed on a gray background (mean luminance ≈100 *cd/m^2^*) to map eccentricity and polar angle. The total run time for both eccentricity and polar angle stimuli was 180 s, comprising five seamless stimulus cycles of 32 s duration each along with 4 s and 12 s of task-only periods (no checkerboard stimuli) respectively at the start and the end.

The flickering checkerboard stimuli had adjacent patches of pseudo-randomly chosen colors, with pairwise euclidean distances in the *Lab* color space (quantifying relative perceptual differences between any two colors) of at least 40. Each of these colored patches were plaided with a set of radially moving points. To improve the perceived contrast, the points were either black or white depending on the color of the patch on which the points were located. The lifetime of these points was set to 0.4 s, a new point at a random location was initialised after that. With every flicker, the color of the patches changed to its complementary luminance. Simultaneously, the color changed and the direction of movement of the plaided points also reversed.

Eccentricity encoding was implemented by a concentric flickering ring expanding and contracting across the visual field (0.95°of visual angle in width). The ring was not scaled with cortical magnification factor. The concentric ring traveled across the visual field in 16 equal steps, stimulating every location in the visual field for 2 s. After each cycle, the expanding or the contracting rings were replaced by new rings at the center or the periphery respectively.

Polar angle encoding was implemented by a single moving wedge (clockwise and counterclockwise direction). The opening angle of the wedge was 22.5 degrees. Similar to the eccentricity stimuli, every location in the visual field was stimulated for 2 seconds before the wedge was moved to the next position.

#### Center letter reading task

In order to keep the participants’ attention focused and to minimize eye-movements, they performed a reading task. A black circle (radius 0.4°) was presented as a fixation point at the center of the screen, superimposed on the main stimulus. Within this circle, a randomly selected excerpt of song lyrics was shown as a stream of single letters (0.5° height, letter frequency 1.5 Hz, 85% duty cycle) throughout the entire length of a run. Participants had to fixate, as they were unable to perform the reading task otherwise. After each acquisition run, participants were presented with a question related to the previously read text. They were given two probable answers, to which they replied by corresponding button press (index or middle finger of their right hand). These question only served the purpose of keep participants attentive — and were otherwise irrelevant. The correctness of the responses was not evaluated.

#### Procedure

Participants performed four acquisition runs in a single session with a total duration of 12 min, with short breaks in-between and without moving out of the scanner. In each run, participants performed the center reading task while passively watching the contracting, counter-clockwise rotating, expanding, and clockwise rotating stimuli in exactly this sequential order. For the retinotopic mapping experiment, 90 volumes of fMRI data were acquired for each run.

### Localizer for higher visual areas

#### Stimulus

All the stimuli for this experiment were used in a previous study^19^. There were 24 unique grayscale images from each of six stimulus categories: human faces, human bodies without heads, small objects, houses and outdoor scenes comprising of nature and street scenes, and phase scrambled images (Figure 1B). Mirrored views of these 24x6 images were also used as stimuli. The original images were converted to grayscale and scaled to a resolution of 400x400px. Images were matched in luminance using `lumMatch` in the SHINE toolbox^20^ to a mean and standard deviation of 128 and 70 respectively. The original images of human faces and houses were produced in the Haxby lab at Dartmouth College; human body images were obtained from Paul Downing’s lab at Bangor University^11,21^; images of small objects were obtained from the Bank of Standardized stimuli (BOSS)^22,23^; outdoor natural scenes are a collection of personal images and public domain resources; and street scenes are taken from the CBCL Street scene database^24^. Stimulus images were displayed at a size of approximately 10°x10° of visual angle.

**Figure 1.**
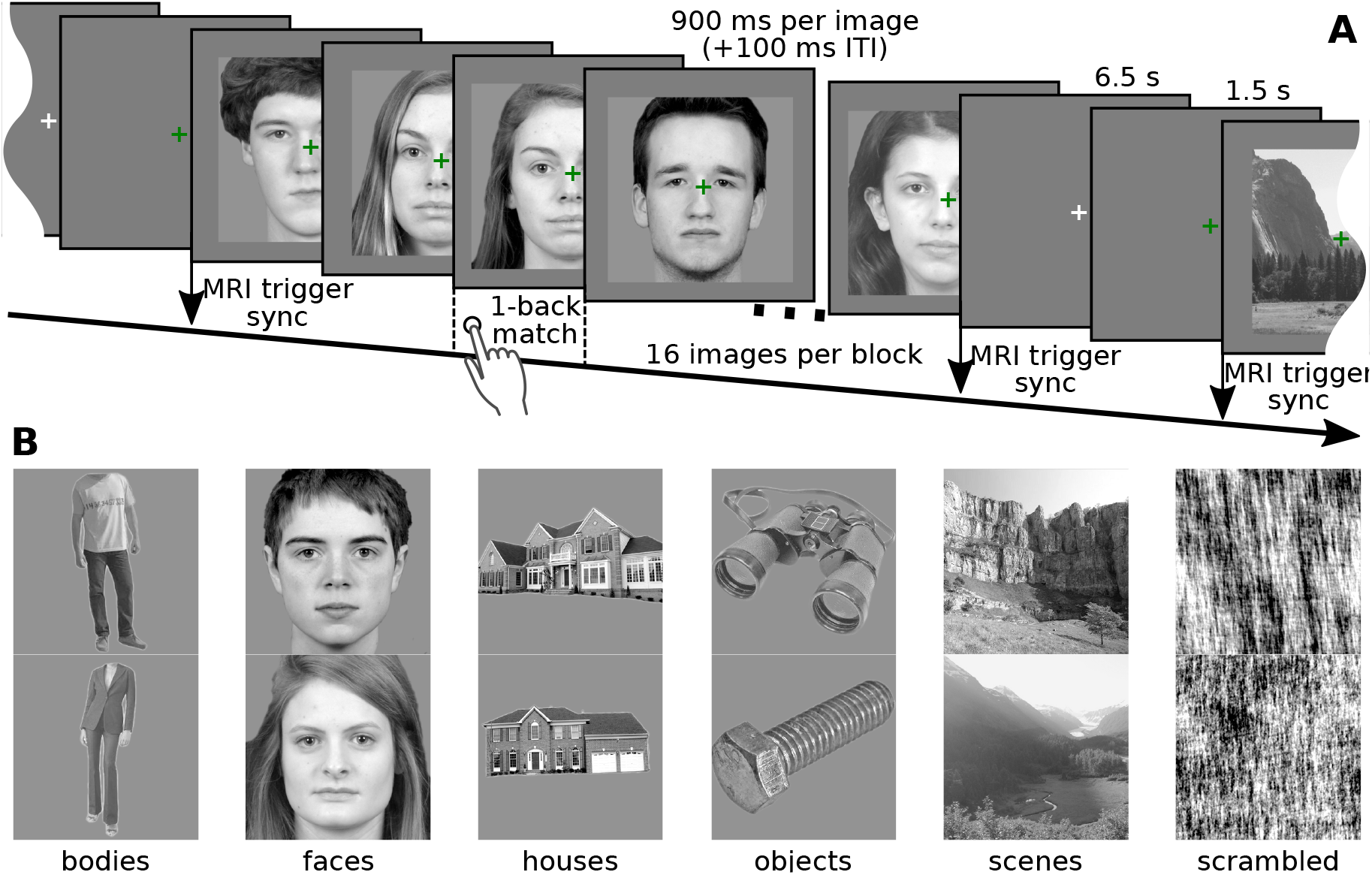
Experiment design. (A) In each block, 16 unique images were presented on a medium-gray background (with a superimposed green fixation cross). Each image was shown for 900ms and images were separated in time by 100 ms. The participant’s task was to press a button (index finger, right hand) when any image was immediately followed by its mirrored equivalent. These events happened randomly either once, twice, or never in each block. In order to alert the participant, the fixation cross turned green 1.5 s prior to the start of a block, remained green throughout a block, and was white during the rest period. The start of each block was synchronized with the MR volume acquisition trigger pulse. Stimulus blocks were separated by 8s of fixation. (B) Example images for all six stimulus categories.

### Procedure

Participants were presented with four block-design runs, with two 16 s blocks per stimulus category in each run, while they also performed a one-back matching task to keep them attentive. The order of blocks was randomized so that all six conditions appeared in random order in both the first and second halves of a run. However, due to a coding error, the block-order was identical across all four runs; though the actual per-block image sequence was a different random sequence for each run. The block configuration and implementation of the matching task are depicted in Figure 1A. 156 fMRI volumes were acquired during each experiment run.

### Movie frame localizer

A third stimulation paradigm was implemented to collect BOLD fMRI data for an independent localization of voxels that show a response to basic visual stimulation in areas of the visual field covered by the movie stimulus used in the companion article^7^. The stimulus was highly similar to the one used for the retinotopic mapping, but instead of isolated rings and wedges, the dynamic stimulus covered either the full rectangle of the movie frame (without the horizontal bars at the top and bottom) or just the horizontal bars. The stimulus alternated every 12 s, starting with the movie frame rectangle. A total of four stimulus alternation cycles were presented — starting synchronized with the acquisition of the first fMRI volume. A total of 48 volumes were acquired. During stimulation, participants performed the same reading task as in the retinotopic mapping session, hence a localization of responsive voxels assumes a central fixation and can only be considered as an approximation of the responsive area of the visual cortex during the movie session, where eye movements were permitted.

### Code availability

All custom source code for data conversion from raw, vendor-specific formats into the de-identified released form is included in the data release (`code/rawdata conversion`). fMRI data conversion from DICOM to NIfTI format was performed with heudiconv (https://github.com/nipy/heudiconv), and the de-identification of these images was implemented with mridefacer (https://github.com/hanke/mridefacer).

The data release also contains the implementations of the stimulation paradigms in code/stimulus/. Moreover, analysis code for visual area localization and retinotopic mapping is available in two dedicated repositories at github.com/psychoinformatics-de/studyforrest-data-visualrois, and github.com/psychoinformatics-de/studyforrest-data-retinotopy, respectively.

## Data Records

This dataset is compliant with the Brain Imaging Data Structure (BIDS) specification^25^, which is a new standard to organize and describe neuroimaging and behavioral data in an intuitive and common manner. Extensive documentation of this standard is available at http://bids.neuroimaging.io. This section provides information about the released data, but limits its description to aspects that extends the BIDS specifications. For a general description of the dataset layout and file naming conventions, the reader is referred to the BIDS documentation. In summary, all files related to the data acquisitions for a particular participant described in this manuscript can be located in a `sub-<ID>/ses-localizer/` directory, where ID is the numeric subject code.

All data records listed in this section are available on the OpenfMRI portal (dataset accession number: `ds000113d`) at http://openfmri.org/dataset/ds000113d as well as on Github/ZENODO^26^.

In order to de-identify data, information on center-specific study and subject codes have been removed using an automated procedure. All human participants were given sequential integer IDs. Furthermore, all BOLD images were “de-faced” by applying a mask image that zeroed out all voxels in the vicinity of the facial surface, teeth, and auricles. For each image modality, this mask was aligned and re-sliced separately. The resulting tailored mask images are provided as part of the data release to indicate which parts of the image were modified by the de-facing procedure (de-face masks carry a `_defacemask` suffix to the base file name).

In addition to the acquired primary data described in this section, we provide results of validation analysis described below. These are: 1) manually titrated ROI masks for visual areas localized for all participants (github.com/psychoinformatics-de/studyforrest-data-visualrois); and 2) volumetric and surface-projected eccentricity and polar angles maps from retinotopic mapping analysis (github.com/psychoinformatics-de/studyforrest-data-retinotopy).

### fMRI data

Each image time series in NIfTI format is accompanied by a JSON sidecar file that contains a dump of the original DICOM metadata for the respective file. Additional standardized metadata is available in the task-specific JSON files defined by the BIDS standard.

### Retinotopic mapping

fMRI data files for the retinotopic mapping contain a `*ses-localizer_task-retmap*` bold in their file name. Specifically the `retmapclw, retmapccw, retmapcexp`, and `retmapcon` file name labels respecitvely indicate stimulation runs with clockwise and counterclockwise rotating wedges, and expanding and contracting rings.

### Higher visual area localizers

fMRI data files for the visual area localizers contain a `*ses-localizer_task-objectcategories*_bold` in their file name. The stimulation timing for each acquisition run is provided in corresponding `*_events.tsv` files. These three-column text files describe the onset and duration of stimulus block and identify the associated stimulus category (`trial_type`).

### Movie frame localizer

fMRI data files for the movie frame localizer contain a `*ses-localizer_task-movielocalizer*_bold` in their file name.

### Physiological recordings

Time series of pleth pulse and respiratory trace are provided for all BOLD fMRI scans in a compressed three-column text file: volume acquisition trigger, pleth pulse, and respiratory trace (file name scheme: `_recording-cardresp_physio.tsv.gz`). The scanner’s built-in recording equipment does not log the volume acquisition trigger nor does it record a reliable marker of the acquisition start. Consequently, the trigger log has been reconstructed based on the temporal position of a scan’s end-marker, the number of volumes acquired, and under the assumption of an exactly identical acquisition time for all volumes. The time series have been truncated to start with the first trigger and end after the last volume has been acquired.

## Technical Validation

All image analyses presented in this section were performed on the released data in order to test for negative effects of the de-identification procedure on subsequent analysis steps.

During data acquisition, (technical) problems were noted in a log. All known anomalies and their impact on the dataset are detailed in Table 1.

**Table 1.** Overview of known data anomalies (F: functional data, P: physiological recordings during fMRI session) for all paradigms (RM: retinotopic mapping, VL: visual localizer, MV: movie localizer).

| Modality | Paradigm | Participant | Run | Description |
| --- | --- | --- | --- | --- |
| F |  | 16 | 4 | excessive motion (rotation) |
| F |  | 10 | 3–4 | excessive motion (translation) |
| F | VL | 20 | 1–4 | excessive motion from coughing |
| F | RM | 20 | 1–4 | reported discomfort during scan |
| P | MV,VL,RM | 2 | all | no data have been acquired |

### Temporal signal-to-noise ratio (tSNR)

Data acquisition was executed using the R5 software version of the scanner vendor. With this version, the vendor changed the frequency of the Spectral Presaturation by Inversion Recovery (SPIR) pulse from the previously 135Hz to 220 Hz in order to increase fat suppression efficiency. After completion of data acquisition, it was discovered that the new configuration led to undesired interactions with pulsations in the cerebrospinal fluid in the ventricles, which resulted in a reduced temporal stability of the MR signal around the ventricles. Figure 2A illustrates the magnitude and spatial extent of this effect. Despite this issue, the majority of voxels show a tSNR of ≈70 or above (Figure 2B), as can be expected with a voxel volume of about 27 mm^3^ and with 3 Tesla acquisition^27^.

**Figure 2.**
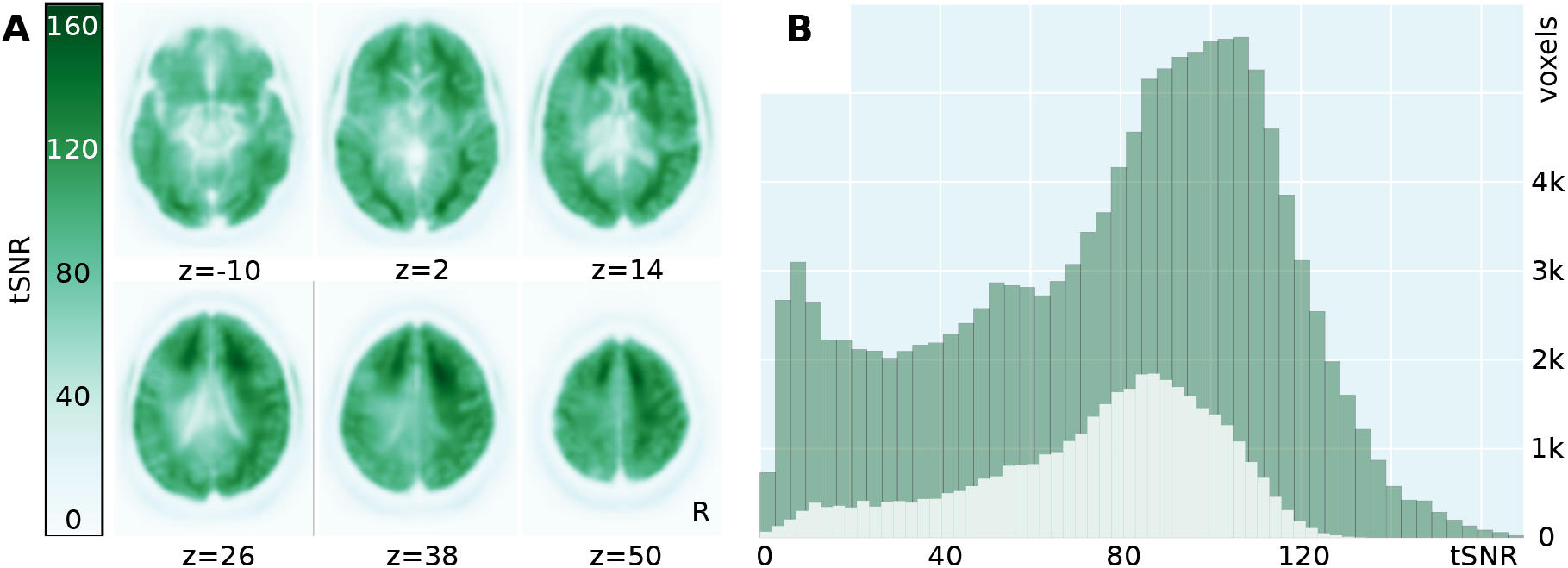
Average temporal signal-to-noise ratio (tSNR) across all acquisitions, including the fMRI data described in the companion article^7^. tSNR was computed independently from motion-corrected and linearly detrended BOLD fMRI time series for each scan. The resulting statistics were projected into group space for averaging across scans and participants (n=255). (A) Spatial distribution of average tSNR across the brain. In the vicinity of the ventricles, tSNR is reduced due to a suboptimal fat suppression SPIR pulse frequency. This artifact amplifies the expected U-shape of the spatial SNR profile of a 32channel head coil. (B) Histograms of average tSNR scores. The dark shaded histogram shows the tSNR distribution of all voxels in an approximate brain mask (MNI template brain mask); the lighter shaded histogram shows the tSNR distribution of 20% of voxels with the largest probability of sampling gray matter, as indicated by FSL’s gray matter prior volume (`avg152T1_gray.nii.gz`) for the MNI template image.

### Retinotopic mapping analysis

Many regions of interest (ROI) in the human visual system follow a retinotopic organization^16,17,28^. The primary areas like V1 and V2 are also provided as labels with the Freesurfer segmentation using the `recon-all` pipeline^29^. But the higher visual areas (V3, VO, PHC, etc) need to be localized by retinotopic mapping^30–33^ or probability maps^34,35^.

We implemented a standard analysis pipeline for the acquired fMRI data based on standard algorithms publicly available in the software packages Freesurfer^29^, FSL^36^, and AFNI^37^. All analysis steps were performed on a computer running the (Neuro)Debian operating system^15^, and all necessary software packages (except for Freesurfer) were obtained from system software package repositories.

BOLD images time series for all scans of the retinotopic mapping paradigm were brain-extracted using FSL’s BET and aligned (rigid-body transformation) to a participant-specific BOLD template image. All volumetric analysis was performed in this image space. An additional rigid-body transformation was computed to align the BOLD template image to the previously published cortical surface reconstructions based on T1 and T2-weighted structural images of the respective participants^1^ for later delineation of visual areas on the cortical surface. Using AFNI tools, time series images were also “deobliqued” (`3dWarp`), slice time corrected (`3dTshift`), and temporally bandpass-filtered (`3dBandpass` cutoff frequencies set to 0.667/32 Hz and 2/32 Hz, where 32 s is the period of both the ring and the wedge stimulus).

For angle map estimation, AFNI’s `waver` command was used to create an ideal response time series waveform based on the design of the stimulus. The bandpass filtered BOLD images were then processed by the `3dRetinoPhase` (`DELAY` phase estimation method was based on the response time series model). Expanding and contracting rings, as well as clockwise and counter-clockwise wedge stimuli, were jointly used to generate average volumetric phase maps representing eccentricity and polar angles for each participant. Polar angle maps were adjusted for a shift in the starting position of the wedge stimulus compared between the two rotation directions. The phase angle representations, relative to the visual field, are shown in Figure 3A. As an overall indicator of mapping quality, Figure 3B shows the distribution of the polar angle representations across all voxels in the MNI occipital lobe mask combined for all participants.

For visualization and subsequent delineation, all volumetric angle maps (after correction) were projected onto the cortical surface mesh of the respective participant using Freesurfer’s `mri_vol2surf` command — separately for each hemisphere. In order to illustrate the quality of the angle maps, the subjectively best, average, and worst participants (respectively: participant 1, 10, and 9) have been selected on the basis of visual inspection. Figure 3C shows the eccentricity maps on the left panel and the polar angle maps for both hemispheres on the right panel. A table summarizing the results of the manual inspections of all surface maps is available at github.com/psychoinformatics-de/studyforrest-data-retinotopy/qa. Delineations of the visual areas depicted in Figure 3C were derived according to Kaule et al. (page 4)^38^. Further details on the procedure can be found in^31,32,39^.

**Figure 3.**
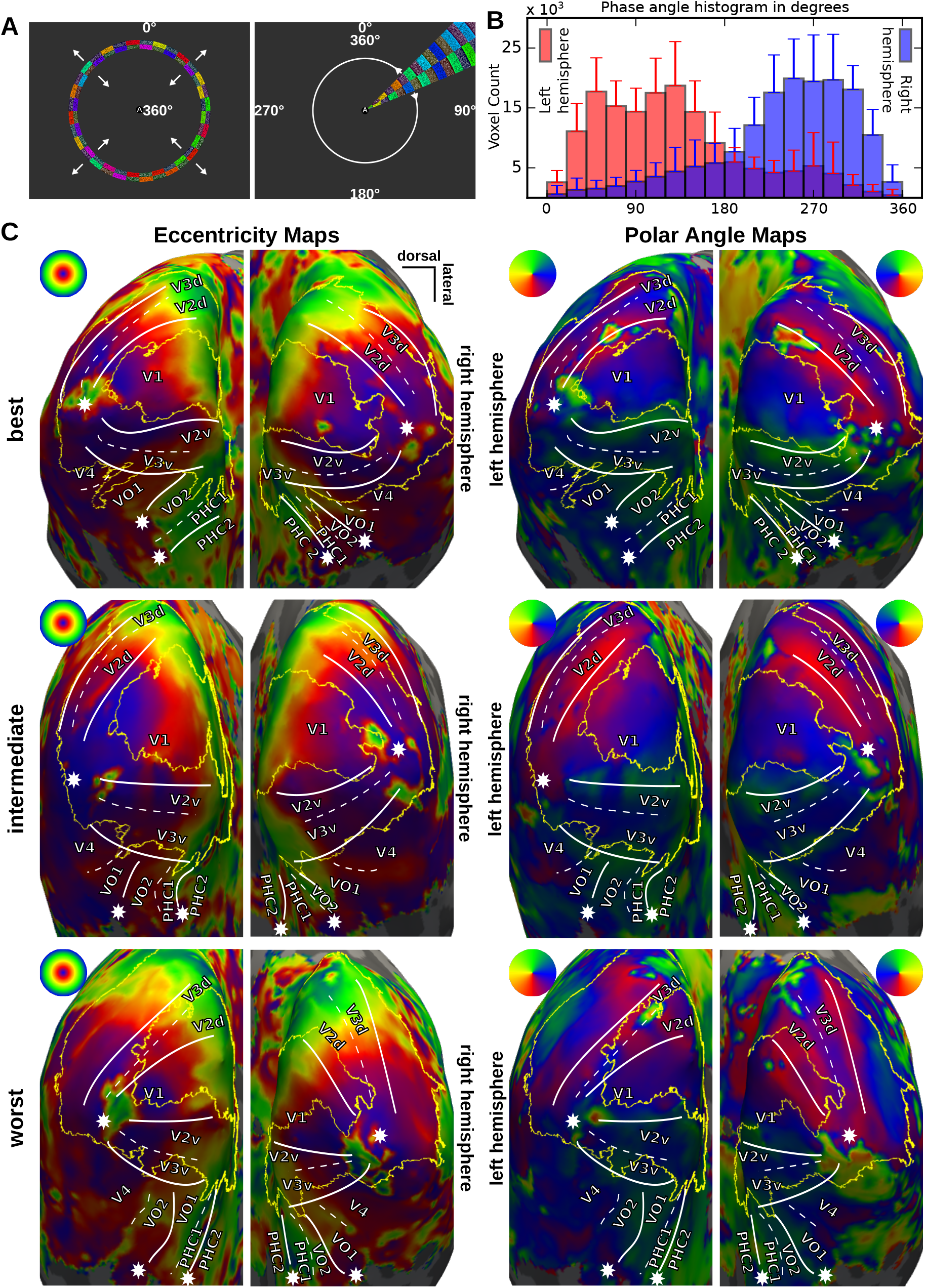
(A) Ring and wedge stimuli with continuous central letter reading task to encourage fixation. White numbers indicate the respective phase angle encoding. (B) Histogram of polar angles for all voxels in the MNI occipital lobe mask for the left and right hemisphere. Error bars indicate standard deviation across all subjects. (C) Inflated occipital cortex surface maps for eccentricity and polar angle for the best, intermediate, and worst participants: participants 1, 10, and 9 respectively. White lines indicate manually delineated visual area boundaries; stars mark the center of the visual field; yellow lines depict the outline of the autogenerated Freesurfer V2 label^1^ for comparison. All maps are constrained to the MNI occipital lobe mask.

### Localization of higher visual areas

To localize higher visual areas for each participant, we implemented a standard two-level general linear model (GLM) analysis using the FEAT component in FSL. BOLD image time series were slice-time-corrected, masked with a conservative brain mask, spatially smoothed (Gaussian kernel, 4 mm FWHM), and temporally high-pass filtered using a cutoff period of 100 s. For each acquisition run, we defined the stimulation design using six boxcar functions, one for each condition (bodies, faces, houses, small objects, landscapes, scrambled images), such that each stimulation block was represented as a single contiguous 16 s segment. The GLM design, comprised of these six regressors, convolved with FSL’s “Double-Gamma HRF” as a model hemodynamic response function model. Temporal derivatives of those regressors were also included in the design matrix, and it was subjected to the same temporal filtering as the BOLD time series.

At the first level, we defined a series of *t*-contrasts to implement different localization procedures found in the literature^40^. The *strict* set included one contrast per target region of interest and involved all stimulus conditions (one condition vs. all others, except for the PPA contrast, where houses/landscapes were contrasted with all other conditions). The relaxed set included structurally similar contrasts as the strict set, but the number of contrast condition was reduced, for example: the FFA contrast was defined as faces vs. small objects and scrambled images. Lastly, the *simple* set contained only contrasts of one (*e*.*g*., faces) or two related conditions (*e*.*g*., houses and landscapes) against responses to scrambled images.

The GLM analysis was performed for each experiment run individually, and afterwards results were aggregated in a within-subject second-level analysis by averaging. Statistical evaluation (fixed-effects analysis) and cluster-level thresholding were performed at the second level using a cluster forming threshold of *Z* >1.64 and a corrected cluster probability threshold of *p* <0.05.

We defined category-selective regions starting with the contrast clusters that survived second-level analysis for each participant. For each region of interest, we started with the most conservative contrast (strict set) by using a threshold of *t* = 2.5 and looked for clusters with at least 20 voxels (using AFNI). We titrated the threshold in the range of [2,3] until we found an isolated cluster for the localizer region of interest. If a cluster was not found or not isolated, we used a contrast from the relaxed set, or finally the simple set, and repeated the process until we found a cluster that matched the expected anatomical location based on literature for FFA/OFA41, PPA9, LOC12, and EBA11.

Figure 4 depicts the results of this procedure for all regions of interest by means of localization overlap across all participants on the cortical surface of the MNI152 brain. Detailed participant-specific information is provided in Table 2 (face-responsive regions), Table 3 (scene and place responsive regions), and Table 4 (early visual areas and LOC). Both the spatial localization of regions in the groups of participants, as well as the frequency of localization success, approximately matches reports in the literature (for example8).

**Figure 4.**
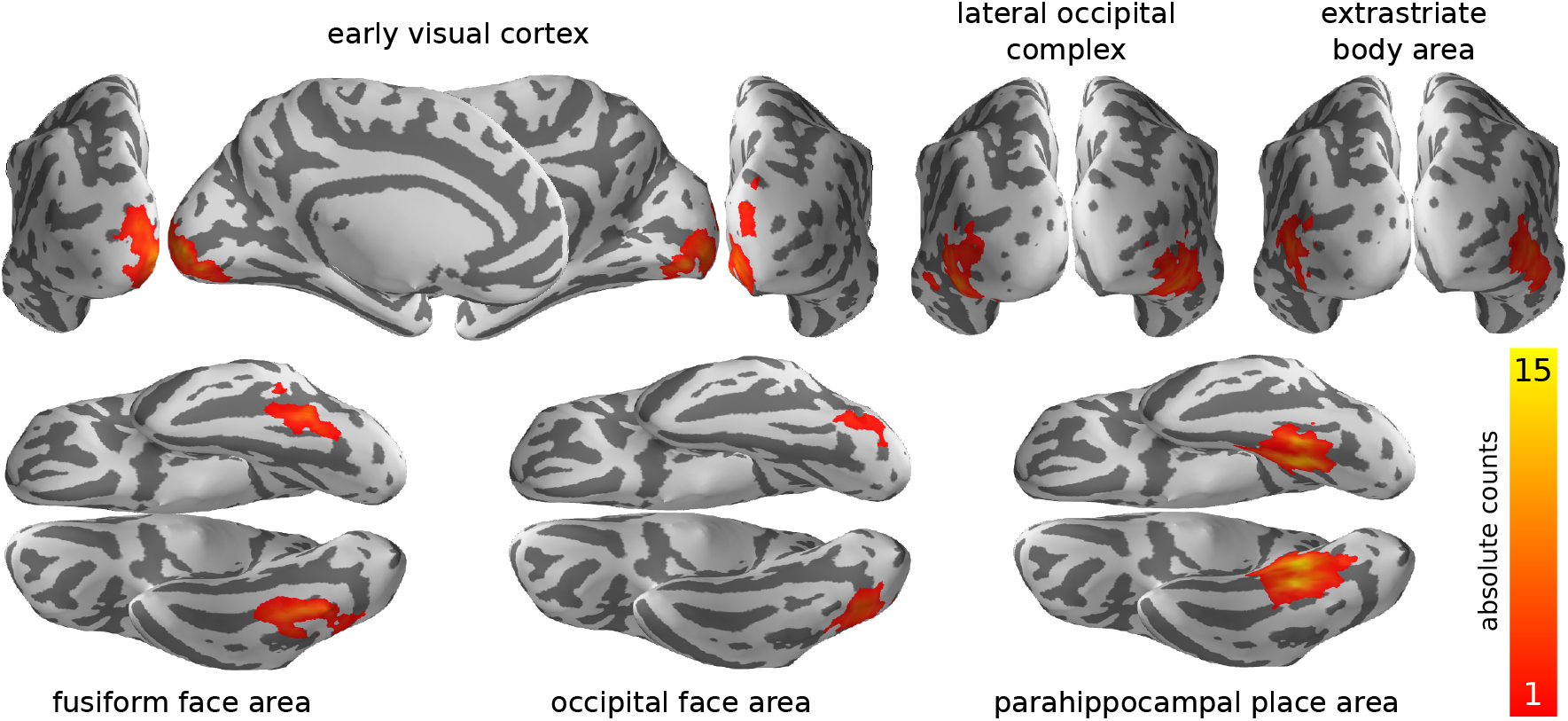
Spatial overlap of individually located regions of interest (ROI). Individual ROI masks were projected into MNI space and summed. Absolute value are shown for nodes on the reconstructed surface of the MNI152 brain. The magnitude is affected by both the spatial variability of ROIs across brains and the localization failures for individual ROIs and participants. See Tables 2, 3, and 4 for individual results.

**Table 2.** Individual localization results after manual titration for face-responsive regions. All statistics correspond to 2nd-level Z-scores; all coordinates are in MNI-space millimeters; ROI volume is reported in cm3; the voxel count corresponds to 2.5 mm isotropic voxels of a participants-specific template image. The last column indicates the underlying contrast type of the statistics maps an ROI definition was based on. Each row corresponds to a single isolated cluster. Multiple rows per participants and ROI indicated the presence of multiple isolated clusters.

|  |  |  |  |  |  |  | max location (MNI) |  |  | center of mass (MNI) |  |  |  |
| --- | --- | --- | --- | --- | --- | --- | --- | --- | --- | --- | --- | --- | --- |
| subj. | min | mean | med. | max | vol. | vox. | X | Y | Z | X | Y | Z | c. type |
| Right fusiform face area (rFFA) |  |  |  |  |  |  |  |  |  |  |  |  |  |
| 1 | 2.50 | 2.83 | 2.77 | 3.59 | 1.08 | 69 | 46.3 | -48.1 | -25.6 | 39.1 | -45.0 | -23.6 | strict |
| 1 | 2.50 | 2.97 | 2.92 | 3.75 | 1.23 | 79 | 48.8 | -58.0 | -21.6 | 42.7 | -59.2 | -18.7 | strict |
| 2 | 2.50 | 3.03 | 2.98 | 3.80 | 2.02 | 129 | 48.8 | -73.1 | -13.1 | 42.5 | -65.4 | -12.2 | strict |
| 3 | 2.24 | 2.52 | 2.44 | 3.10 | 0.95 | 61 | 56.0 | -50.5 | -25.8 | 46.2 | -46.4 | -25.0 | relaxed |
| 4 | 2.00 | 2.45 | 2.38 | 3.37 | 1.20 | 77 | 46.4 | -78.1 | -11.0 | 40.2 | -59.4 | -15.1 | simple |
| 5 | 2.00 | 2.60 | 2.57 | 3.30 | 1.75 | 112 | 43.9 | -53.1 | -23.6 | 39.4 | -48.4 | -21.7 | strict |
| 9 | 2.60 | 2.95 | 2.91 | 3.56 | 1.08 | 69 | 41.5 | -67.7 | -22.1 | 37.2 | -63.1 | -14.7 | strict |
| 9 | 2.60 | 3.05 | 3.03 | 3.63 | 0.69 | 44 | 46.3 | -45.8 | -23.2 | 41.7 | -44.6 | -22.9 | strict |
| 10 | 2.66 | 3.11 | 3.15 | 3.67 | 1.02 | 65 | 53.6 | -48.2 | -23.3 | 44.4 | -48.8 | -18.0 | strict |
| 14 | 2.60 | 2.80 | 2.76 | 3.23 | 1.25 | 80 | 53.6 | -40.7 | -27.6 | 40.0 | -53.1 | -19.8 | simple |
| 15 | 2.75 | 3.03 | 3.01 | 3.46 | 1.55 | 99 | 56.0 | -68.1 | -15.2 | 46.1 | -70.2 | -9.7 | strict |
| 16 | 2.50 | 3.03 | 3.00 | 3.72 | 1.95 | 125 | 46.3 | -53.7 | -14.3 | 40.4 | -50.7 | -14.5 | simple |
| 17 | 2.50 | 2.90 | 2.88 | 3.51 | 1.78 | 114 | 43.8 | -36.4 | -20.3 | 35.5 | -42.5 | -22.5 | simple |
| 19 | 2.90 | 3.24 | 3.20 | 3.78 | 1.09 | 70 | 46.3 | -60.3 | -24.0 | 40.2 | -57.7 | -17.5 | simple |
| 20 | 2.50 | 2.83 | 2.82 | 3.42 | 1.08 | 69 | 48.7 | -48.4 | -21.0 | 41.6 | -49.8 | -19.5 | strict |
| Left fusiform face area (lFFA) |  |  |  |  |  |  |  |  |  |  |  |  |  |
| 1 | 2.50 | 2.71 | 2.70 | 3.06 | 0.27 | 17 | -33.6 | -41.0 | -27.3 | -38.7 | -43.2 | -27.3 | strict |
| 1 | 2.50 | 2.72 | 2.67 | 3.13 | 0.31 | 20 | -36.0 | -58.1 | -23.6 | -42.1 | -59.0 | -23.6 | strict |
| 2 | 2.50 | 3.02 | 3.01 | 3.70 | 1.06 | 68 | -36.0 | -68.4 | -14.9 | -39.6 | -65.4 | -15.3 | strict |
| 5 | 2.00 | 2.44 | 2.44 | 3.22 | 0.94 | 60 | -38.5 | -48.8 | -18.4 | -43.2 | -52.9 | -21.6 | strict |
| 6 | 2.50 | 2.79 | 2.80 | 3.23 | 1.06 | 68 | -33.6 | -68.2 | -17.2 | -39.1 | -61.0 | -15.0 | simple |
| 9 | 2.60 | 2.89 | 2.89 | 3.58 | 1.20 | 77 | -26.3 | -71.1 | -10.4 | -33.0 | -70.8 | -15.1 | strict |
| 10 | 2.66 | 3.09 | 3.09 | 3.67 | 0.67 | 43 | -36.0 | -61.0 | -16.8 | -42.7 | -53.5 | -18.8 | strict |
| 14 | 2.60 | 2.98 | 2.93 | 3.54 | 1.06 | 68 | -38.4 | -60.8 | -19.1 | -43.3 | -53.3 | -22.0 | simple |
| 15 | 2.25 | 2.52 | 2.48 | 3.06 | 1.31 | 84 | -36.0 | -53.2 | -25.6 | -43.7 | -48.3 | -22.9 | simple |
| 16 | 2.50 | 2.82 | 2.80 | 3.32 | 1.11 | 71 | -36.0 | -66.1 | -12.4 | -41.1 | -53.1 | -17.0 | simple |
| 17 | 2.50 | 2.79 | 2.75 | 3.32 | 1.06 | 68 | -33.6 | -51.0 | -20.9 | -37.6 | -42.6 | -17.6 | simple |
| 19 | 2.72 | 2.97 | 2.97 | 3.28 | 0.47 | 30 | -36.0 | -66.0 | -14.7 | -39.2 | -59.0 | -15.0 | relaxed |
| 20 | 2.50 | 2.78 | 2.75 | 3.25 | 0.62 | 40 | -31.2 | -53.9 | -14.1 | -37.6 | -55.6 | -16.4 | strict |
| Right occipital face area (rOFA) |  |  |  |  |  |  |  |  |  |  |  |  |  |
| 3 | 2.50 | 2.72 | 2.68 | 3.17 | 0.98 | 63 | 56.0 | -65.7 | -15.0 | 47.1 | -72.8 | -13.4 | relaxed |
| 4 | 2.00 | 2.14 | 2.13 | 2.35 | 0.70 | 45 | 34.3 | -90.4 | -7.0 | 28.8 | -87.3 | -4.3 | simple |
| 5 | 2.00 | 2.23 | 2.16 | 2.79 | 0.48 | 31 | 46.4 | -85.3 | -11.4 | 37.9 | -80.8 | -15.2 | relaxed |
| 9 | 2.60 | 3.01 | 3.02 | 3.68 | 1.08 | 69 | 48.8 | -75.3 | -15.5 | 41.6 | -79.9 | -13.5 | strict |
| 10 | 2.66 | 3.01 | 3.02 | 3.47 | 0.69 | 44 | 46.4 | -73.2 | -10.8 | 41.3 | -73.3 | -9.4 | strict |
| 14 | 2.60 | 2.77 | 2.75 | 3.06 | 1.03 | 66 | 56.0 | -71.4 | -1.4 | 42.7 | -79.3 | -6.4 | simple |
| 15 | 2.75 | 3.06 | 2.99 | 3.71 | 1.53 | 98 | 46.3 | -48.6 | -18.7 | 40.7 | -50.1 | -16.4 | strict |
| 16 | 2.50 | 2.87 | 2.85 | 3.56 | 2.23 | 143 | 53.6 | -78.8 | 0.6 | 41.8 | -78.8 | -9.3 | simple |
| 17 | 2.40 | 2.68 | 2.64 | 3.31 | 0.86 | 55 | 51.2 | -67.9 | -17.5 | 39.4 | -73.9 | -16.6 | strict |
| 19 | 2.90 | 3.18 | 3.12 | 3.69 | 0.92 | 59 | 46.4 | -83.3 | -4.3 | 41.6 | -79.3 | -12.2 | simple |
| 20 | 2.45 | 2.82 | 2.82 | 3.31 | 0.53 | 34 | 46.4 | -80.9 | -4.2 | 41.3 | -77.6 | -8.1 | strict |
| Left occipital face area (lOFA) |  |  |  |  |  |  |  |  |  |  |  |  |  |
| 10 | 2.66 | 3.01 | 2.97 | 3.48 | 0.66 | 42 | -26.3 | -78.0 | -15.4 | -33.9 | -83.2 | -12.1 | strict |
| 14 | 2.60 | 2.83 | 2.79 | 3.19 | 0.48 | 31 | -31.1 | -89.9 | -18.4 | -37.5 | -85.8 | -15.0 | simple |
| 15 | 3.00 | 3.25 | 3.23 | 3.55 | 0.81 | 52 | -35.9 | -87.9 | -11.3 | -41.9 | -75.6 | -14.8 | simple |
| 17 | 2.20 | 2.51 | 2.46 | 2.99 | 0.67 | 43 | -35.9 | -82.3 | -22.6 | -38.5 | -78.9 | -19.5 | simple |
| 19 | 3.00 | 3.24 | 3.22 | 3.65 | 0.64 | 41 | -35.9 | -85.4 | -13.5 | -39.6 | -75.2 | -15.2 | simple |
| 20 | 2.45 | 2.70 | 2.64 | 3.23 | 0.91 | 58 | -28.7 | -82.0 | -27.3 | -38.4 | -79.5 | -19.8 | strict |

**Table 3.**
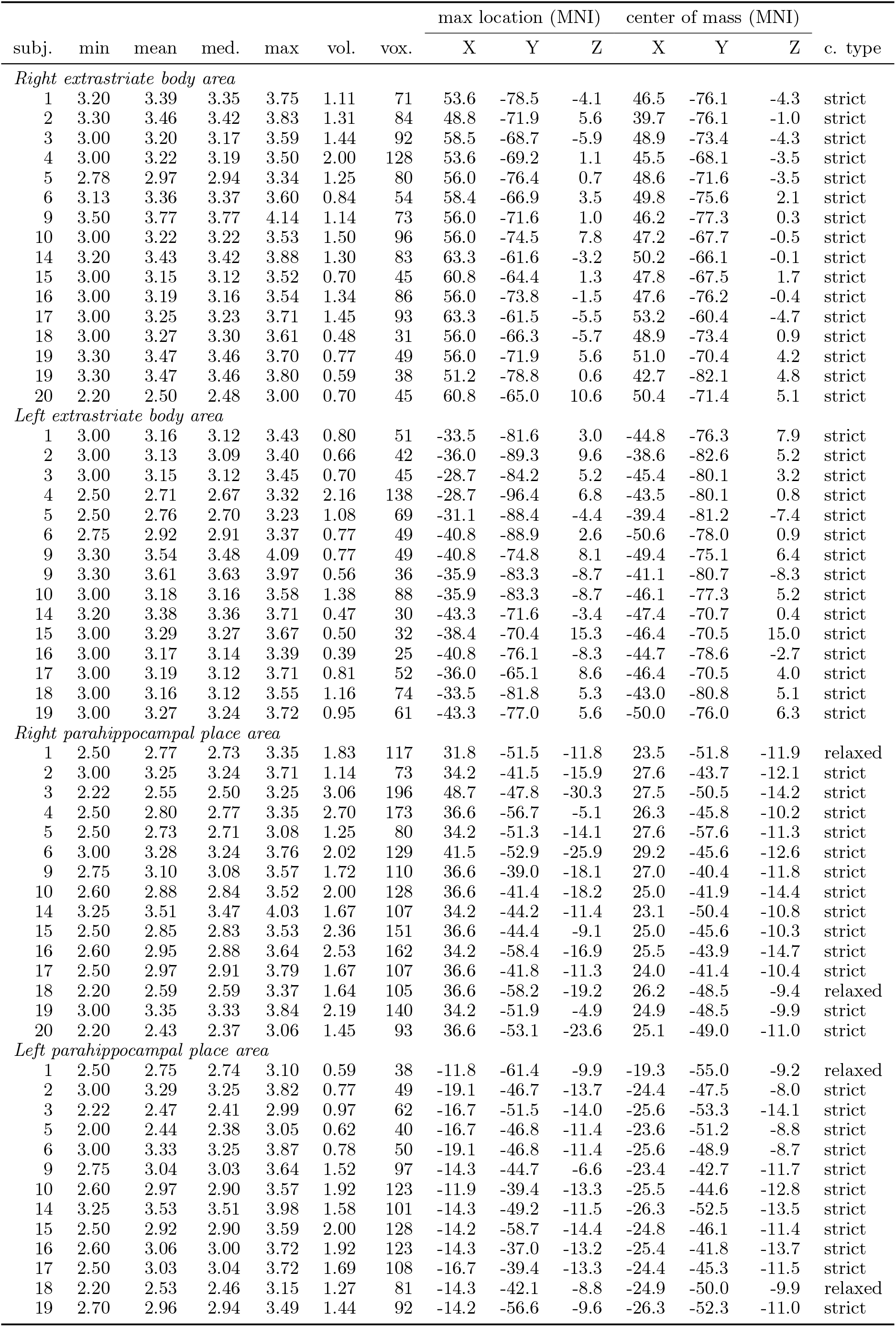
Individual localization results after manual titration for body- and place-responsive regions. Table semantics are identical to Table 2.

**Table 4.** Individual localization results after manual titration for early visual cortex and lateral occipital complex. Table semantics are identical to Table 2.

| subj. | min | mean | med. | max | vol. | vox. | max location (MNI) |  |  | center of mass (MNI) |  |  | c. type |
| --- | --- | --- | --- | --- | --- | --- | --- | --- | --- | --- | --- | --- | --- |
|  |  |  |  |  |  |  | X | Y | Z | X | Y | Z |  |
| Right lateral occipital complex |  |  |  |  |  |  |  |  |  |  |  |  |  |
| 1 | 2.32 | 2.53 | 2.52 | 2.92 | 1.61 | 103 | 56.0 | -66.3 | -5.7 | 43.5 | -80.9 | -9.7 | simple |
| 2 | 2.50 | 2.75 | 2.73 | 3.24 | 3.20 | 205 | 53.6 | -71.4 | -1.4 | 39.1 | -82.0 | -4.8 | simple |
| 3 | 2.50 | 2.75 | 2.71 | 3.24 | 2.34 | 150 | 58.5 | -68.1 | -15.2 | 49.5 | -67.3 | -14.7 | simple |
| 4 | 2.50 | 2.65 | 2.65 | 2.82 | 0.83 | 53 | 34.3 | -86.1 | 0.2 | 29.5 | -84.3 | 1.6 | simple |
| 5 | 2.40 | 2.62 | 2.59 | 2.94 | 0.75 | 48 | 56.1 | -73.4 | -8.5 | 46.5 | -73.5 | -10.1 | simple |
| 9 | 2.50 | 2.74 | 2.68 | 3.14 | 0.77 | 49 | 41.6 | -85.1 | -13.7 | 36.4 | -83.8 | -1.9 | simple |
| 10 | 2.50 | 2.70 | 2.66 | 3.19 | 0.73 | 47 | 53.6 | -73.8 | -1.5 | 45.8 | -71.8 | -2.8 | simple |
| 14 | 2.50 | 2.68 | 2.67 | 2.99 | 1.86 | 119 | 65.7 | -58.7 | -10.0 | 47.2 | -72.7 | -5.7 | simple |
| 15 | 2.85 | 3.12 | 3.09 | 3.46 | 1.53 | 98 | 56.0 | -68.1 | -15.2 | 46.7 | -69.7 | -6.6 | simple |
| 16 | 2.80 | 3.00 | 2.98 | 3.39 | 2.16 | 138 | 56.0 | -66.3 | -5.7 | 42.9 | -76.4 | -7.8 | simple |
| 17 | 2.65 | 2.88 | 2.86 | 3.34 | 2.44 | 156 | 60.9 | -66.3 | -5.8 | 47.0 | -67.7 | -7.7 | strict |
| 18 | 2.20 | 2.49 | 2.44 | 2.90 | 1.77 | 113 | 56.1 | -73.4 | -8.5 | 43.4 | -77.7 | -7.5 | strict |
| 19 | 2.00 | 2.29 | 2.29 | 2.64 | 1.78 | 114 | 58.4 | -71.9 | 5.6 | 46.6 | -72.5 | 7.6 | simple |
| 20 | 2.00 | 2.24 | 2.18 | 2.82 | 1.08 | 69 | 56.0 | -66.9 | 3.5 | 45.0 | -76.1 | -4.7 | strict |
| Left lateral occipital complex |  |  |  |  |  |  |  |  |  |  |  |  |  |
| 1 | 2.80 | 2.99 | 2.96 | 3.44 | 2.03 | 130 | -33.5 | -90.3 | -11.4 | -41.6 | -78.0 | -12.0 | simple |
| 2 | 2.50 | 2.85 | 2.80 | 3.38 | 2.70 | 173 | -31.1 | -83.4 | -6.4 | -39.7 | -81.3 | -6.0 | simple |
| 3 | 2.50 | 2.72 | 2.67 | 3.30 | 2.20 | 141 | -31.1 | -81.9 | 7.7 | -45.4 | -77.8 | -2.8 | simple |
| 4 | 2.50 | 2.66 | 2.66 | 2.89 | 1.50 | 96 | -31.1 | -91.3 | 2.5 | -40.3 | -80.3 | -2.7 | simple |
| 5 | 2.40 | 2.62 | 2.56 | 3.05 | 1.17 | 75 | -31.1 | -86.3 | 0.4 | -38.6 | -80.8 | -6.5 | simple |
| 9 | 2.50 | 2.83 | 2.78 | 3.35 | 2.22 | 142 | -21.4 | -90.3 | -11.5 | -40.3 | -80.2 | -11.9 | simple |
| 10 | 2.50 | 2.88 | 2.87 | 3.32 | 2.62 | 168 | -33.5 | -83.1 | -11.0 | -42.8 | -79.1 | 1.3 | simple |
| 14 | 2.50 | 2.69 | 2.68 | 3.04 | 1.19 | 76 | -23.8 | -88.1 | -9.0 | -41.4 | -77.5 | -0.3 | simple |
| 14 | 2.50 | 2.75 | 2.69 | 3.29 | 1.64 | 105 | -38.4 | -70.8 | -15.0 | -50.7 | -65.0 | -8.2 | simple |
| 15 | 2.85 | 3.05 | 3.03 | 3.31 | 0.72 | 46 | -36.0 | -78.8 | -3.8 | -48.8 | -74.0 | -1.8 | simple |
| 15 | 2.85 | 3.13 | 3.14 | 3.44 | 0.83 | 53 | -36.0 | -76.1 | -8.3 | -41.8 | -73.2 | -12.4 | simple |
| 16 | 2.80 | 2.95 | 2.94 | 3.20 | 2.12 | 136 | -31.1 | -80.9 | -8.6 | -43.9 | -73.8 | -8.1 | simple |
| 17 | 2.70 | 2.98 | 2.95 | 3.53 | 1.97 | 126 | -35.9 | -77.8 | -17.7 | -47.5 | -70.7 | -9.1 | simple |
| 18 | 2.20 | 2.39 | 2.36 | 2.92 | 1.55 | 99 | -35.9 | -85.4 | -13.5 | -43.4 | -78.8 | -9.1 | strict |
| 19 | 2.00 | 2.37 | 2.37 | 3.02 | 0.55 | 35 | -43.2 | -76.4 | -3.6 | -48.0 | -76.5 | 0.9 | simple |
| 20 | 2.00 | 2.29 | 2.25 | 2.65 | 0.81 | 52 | -31.1 | -93.8 | 4.7 | -36.8 | -88.8 | 4.0 | strict |
| Early visual cortex |  |  |  |  |  |  |  |  |  |  |  |  |  |
| 1 | 2.50 | 2.79 | 2.75 | 3.51 | 3.48 | 223 | 12.5 | -88.3 | -4.5 | -0.3 | -88.1 | -6.4 | strict |
| 2 | 3.00 | 3.21 | 3.17 | 3.58 | 3.22 | 206 | 14.9 | -78.5 | -6.3 | 1.0 | -87.5 | -5.4 | strict |
| 4 | 2.25 | 2.58 | 2.53 | 3.29 | 6.81 | 436 | 19.8 | -90.5 | -7.0 | -1.5 | -92.4 | -4.0 | strict |
| 5 | 2.50 | 2.89 | 2.82 | 3.54 | 4.62 | 296 | 22.2 | -96.5 | 11.3 | 11.4 | -88.7 | -4.3 | strict |
| 5 | 2.50 | 2.90 | 2.85 | 3.53 | 1.50 | 96 | 2.9 | -100.3 | -5.1 | -7.7 | -97.4 | 0.8 | strict |
| 9 | 3.00 | 3.24 | 3.23 | 3.61 | 4.31 | 276 | 17.4 | -90.8 | -2.3 | 3.0 | -90.5 | -4.2 | strict |
| 10 | 2.80 | 3.04 | 3.01 | 3.51 | 4.92 | 315 | 22.2 | -102.5 | -7.7 | 1.8 | -92.2 | 2.1 | strict |
| 14 | 2.50 | 2.81 | 2.76 | 3.29 | 1.12 | 72 | 19.8 | -91.9 | 13.9 | 14.8 | -95.8 | 8.2 | strict |
| 14 | 2.50 | 2.84 | 2.79 | 3.45 | 3.34 | 214 | 7.6 | -83.1 | -8.9 | -5.0 | -92.5 | -1.3 | strict |
| 15 | 2.40 | 2.67 | 2.63 | 3.24 | 1.67 | 107 | 29.4 | -89.8 | 18.7 | 15.4 | -93.3 | 11.9 | strict |
| 15 | 2.40 | 2.79 | 2.77 | 3.38 | 1.53 | 98 | -2.0 | -101.0 | 4.2 | -10.4 | -98.6 | 6.2 | strict |
| 16 | 2.50 | 2.78 | 2.75 | 3.31 | 3.81 | 244 | 22.2 | -96.5 | 11.3 | 4.6 | -91.6 | 0.6 | strict |
| 17 | 2.50 | 2.86 | 2.83 | 3.50 | 3.34 | 214 | 29.4 | -94.7 | 20.7 | 10.3 | -88.5 | 1.5 | strict |
| 17 | 2.50 | 2.87 | 2.89 | 3.44 | 1.86 | 119 | -6.8 | -98.4 | 2.0 | -12.4 | -97.5 | 4.2 | strict |
| 18 | 2.00 | 2.37 | 2.31 | 3.25 | 5.70 | 365 | 27.0 | -75.6 | -13.1 | 2.0 | -88.0 | -5.0 | strict |
| 19 | 3.00 | 3.22 | 3.20 | 3.63 | 1.39 | 89 | 19.8 | -97.7 | -7.4 | 11.1 | -91.4 | -2.7 | strict |
| 19 | 3.00 | 3.22 | 3.21 | 3.60 | 1.56 | 100 | 0.4 | -100.3 | -5.1 | -8.2 | -99.7 | 6.6 | strict |
| 20 | 2.00 | 2.28 | 2.24 | 3.06 | 4.67 | 299 | 22.2 | -91.7 | 11.6 | 0.1 | -93.6 | 0.7 | strict |

## Usage Notes

The procedures we employed in this study resulted in a dataset that is highly suitable for automated processing. Data files are organized according to the BIDS standard^25^. Data are shared in documented standard formats, such as NIfTI or plain text files, to enable further processing in arbitrary analysis environments with no imposed dependencies on proprietary tools. Conversion from the original raw data formats is implemented in publicly accessible scripts; the type and version of employed file format conversion tools are documented. Moreover, all results presented in this section were produced by open source software on a computational cluster running the (Neuro)Debian operating system^15^. This computational environment is freely available to anyone, and it — in conjunction with our analysis scripts — offers a high level of transparency regarding all aspects of the analyses presented herein.

All data are made available under the terms of the Public Domain Dedication and License (PDDL; http://opendatacommons.org/licenses/pddl/1.0/). All source code is released under the terms of the MIT license (http://www.opensource.org/licenses/MIT). In short, this means that anybody is free to download and use this dataset for any purpose as well as to produce and re-share derived data artifacts. While not legally required, we hope that all users of the data will acknowledge the original authors by citing this publication and follow good scientific practise as laid out in the ODC Attribution/Share-Alike Community Norms (http://opendatacommons.org/norms/odc-by-sa/).

## Acknowledgments

We acknowledge the support of the Combinatorial NeuroImaging Core Facility at the Leibniz Institute for Neurobiology in Magdeburg. Michael Hanke was supported by funds from the German federal state of Saxony-Anhalt, Project: Center for Behavioral Brain Sciences. This research was, in part, co-funded by the German Federal Ministry of Education and Research (BMBF 01GQ1112, 01GQ1411) and the US National Science Foundation (NSF 1129855, 1429999) as part of two US-German collaborations in computational neuroscience (CRCNS). We are grateful to the Haxby lab, Paul Downing, the maintainers of the BOSS stimulus library, and the CBCL street scene library for making the stimulus material publicly available. We appreciate the editing efforts of Alex Waite and do publicly acknowledge the excellent beer making traditions of the great state of Wisconsin.

AS contributed to the manuscript and the implementation of stimulation paradigm and validation analysis of the retinotopic mapping. FRK wrote the retinopy analysis, performed retinotopy dataset validation analysis, and contributed to the manuscript. JSG contributed to the implementation of the stimulation paradigm and contributed to the validation analysis of the visual localizers. MBH contributed the examination of visual function and expertise in retinotopic mapping. CH contributed to the visual localizer analysis. JS was the data acquisition lead. MH conducted the study, implemented the stimulation paradigm, performed dataset validation analysis, and wrote the manuscript.

## References

1 Hanke M. et al. A high-resolution 7-Tesla fMRI dataset from complex natural stimulation with an audio movie. Scientific Data 1 (2014). URL http://dx.doi.org/10.1038/sdata.2014.3.

2 Nguyen V. T., Breakspear M., Hu X. & Guo C. C. The integration of the internal and external milieu in the insula during dynamic emotional experiences. NeuroImage 124, 455–463 (2016).

3 Chen, P.-H. C. et al. A reduced-dimension fMRI shared response model. In Cortes, C., Lawrence N. D., Lee D. D., Sugiyama M. & Garnett R. (eds.) Advances in Neural Information Processing Systems 28, 460–468 (Curran Associates, Inc., 2015). URL http://papers.nips.cc/paper/5855-a-reduced-dimension-fmri-shared-response-model.pdf.

4 Hu X., Guo L., Han J. & Liu T. Decoding power-spectral profiles from fmri brain activities during naturalistic auditory experience. Brain Imaging and Behavior 1–11 (2016).

5 Hanke M. et al. High-resolution 7-Tesla fMRI data on the perception of musical genres – an extension to the studyforrest dataset. F1000Research 4:174 (2015).

6 Labs A. et al. Portrayed emotions in the movie “Forrest Gump”. F1000Research 4:92 (2015). URL http://f1000research.com/articles/4-92.

7 Hanke M. et al. Simultaneous fMRI and eye gaze recordings during prolonged natural stimulation – a studyforrest extension. Scientific Data submitted (2016).

8 Kanwisher N., McDermott J. & Chun M. M. The fusiform face area: a module in human extrastriate cortex specialized for face perception. The Journal of Neuroscience 17, 4302–4311 (1997).

9 Epstein R. & Kanwisher N. A cortical representation of the local visual environment. Nature 392, 598–601 (1998).

10 Pitcher D., Walsh V. & Duchaine B. The role of the occipital face area in the cortical face perception network. Experimental Brain Research 209, 481–493 (2011).

11 Downing P. E., Jiang Y., Shuman M. & Kanwisher N. A cortical area selective for visual processing of the human body. Science 293, 2470–2473 (2001).

12 Malach R. et al. Object-related activity revealed by functional magnetic resonance imaging in human occipital cortex. Proceedings of the National Academy of Sciences 92, 8135–8139 (1995).

13 Sengupta A. et al. An extension of the studyforrest dataset for vision research. Scientific Data submitted (2016).

14 Peirce J. PsychoPy–Psychophysics software in Python. Journal of Neuroscience Methods 162, 8–13 (2007).

15 Halchenko Y. O. & Hanke M. Open is not enough. Let’s take the next step: An integrated, community-driven computing platform for neuroscience. Frontier in Neuroinformatics 6 (2012).

16 Engel S. A., Glover G. H. & Wandell B. A. Retinotopic organization in human visual cortex and the spatial precision of functional mri. Cerebral cortex 7, 181–192 (1997).

17 Sereno M. et al. Borders of multiple visual areas in humans revealed by functional magnetic resonance imaging. Science 268, 889–893 (1995).

18 Warnking J. et al. fMRI retinotopic mapping–step by step. NeuroImage 17, 1665–83 (2002).

19 Haxby J. V. et al. A common, high-dimensional model of the representational space in human ventral temporal cortex. Neuron 72, 404–416 (2011). PMC3201764.

20 Willenbockel V. et al. Controlling low-level image properties: the shine toolbox. Behavior Research Methods 42, 671–684 (2010).

21 Downing P. Localizer stimuli for the extrastriate body area. http://pages.bangor.ac.uk/~;pss811/page7/files/EBA_localizer.zip. (2001).

22 Brodeur M., Lepage M. & Gu´erard, K. Bank Of Standardized Stimuli (BOSS). https://sites.google.com/site/bosstimuli/home. Released under CC BY-SA 3.0 (2009).

23 Brodeur M. B., Dionne-Dostie E., Montreuil T. & Lepage M. The Bank of Standardized Stimuli (BOSS), a new set of 480 normative photos of objects to be used as visual stimuli in cognitive research. PloS one 5, e10773 (2010).

24 Bileschi S. CBCL Street scence database. http://cbcl.mit.edu/software-datasets/streetscenes/ (2007).

25 Gorgolewski K. J. et al. The brain imaging data structure: a standard for organizing and describing outputs of neuroimaging experiments. bioRxiv (2016). URL http://biorxiv.org/content/early/2016/02/05/034561. http://biorxiv.org/content/early/2016/02/05/034561.full.pdf.

26 Hanke, M. et al. studyforrest-data-phase2 (2016). URL http://dx.doi.org/10.5281/zenodo.48421.

27 Triantafyllou C. et al. Comparison of physiological noise at 1.5 T, 3 T and 7 T and optimization of fMRI acquisition parameters. NeuroImage 26, 243–250 (2005).

28 Engel S. A. et al. fMRI of human visual cortex. Nature (1994).

29 Dale A. M., Fischl B. & Sereno M. I. Cortical surface-based analysis: I. Segmentation and surface reconstruction. NeuroImage 9, 179–194 (1999).

30 Sereno M. I., Lutti A., Weiskopf N. & Dick F. Mapping the human cortical surface by combining quantitative T1 with retinotopy. Cerebral Cortex 23, 2261–2268 (2012).

31 Arcaro M. J., McMains S. A., Singer B. D. & Kastner S. Retinotopic organization of human ventral visual cortex. The Journal of Neuroscience 29, 10638–10652 (2009).

32 Wandell B. A., Dumoulin S. O. & Brewer A. A. Visual field maps in human cortex. Neuron 56, 366–383 (2007).

33 Silver M. A. & Kastner S. Topographic maps in human frontal and parietal cortex. Trends in Cognitive Sciences 13, 488–495 (2009).

34 Wang L., Mruczek R. E., Arcaro M. J. & Kastner S. Probabilistic maps of visual topography in human cortex. Cerebral Cortex bhu277 (2014).

35 Van Essen, D. C. et al. Mapping visual cortex in monkeys and humans using surface-based atlases. Vision research 41, 1359–1378 (2001).

36 Smith S. et al. Advances in functional and structural MR image analysis and implementation as FSL. NeuroImage 23 Suppl 1, S208–19 (2004).

37 Cox R. W. AFNI: Software for analysis and visualization of functional magnetic resonance neuroimages. Computers and Biomedical Research 29, 162–173 (1996).

38 Kaule F. R. et al. Impact of chiasma opticum malformations on the organization of the human ventral visual cortex. Hum. Brain Mapp. 5093–5105 (2014).

39 Silver M. A. & Kastner S. Topographic maps in human frontal and parietal cortex. Trends in Cognitive Sciences 13, 488–495 (2009).

40 Berman M. G. et al. Evaluating functional localizers: The case of the FFA. NeuroImage 50, 56–71 (2010).

41 Kanwisher N. Functional specificity in the human brain: a window into the functional architecture of the mind. Proceedings of the National Academy of Sciences 107, 11163–11170 (2010).

1 Hanke, M. et al. OpenfMRI (2016).

